# Regulation of urea cycle by reversible high stoichiometry lysine succinylation

**DOI:** 10.1101/2022.06.24.497535

**Authors:** Ran Zhang, Xueshu Xie, Chris Carrico, Jesse G. Meyer, Lei Wei, Joanna Bons, Jacob Rose, Rebeccah Riley, Ryan Kwok, Prasanna A. Kumaar, Wenjuan He, Yuya Nishida, Xiaojing Liu, Jason W. Locasale, Birgit Schilling, Eric Verdin

## Abstract

The post-translational modification, lysine succinylation is implicated in the regulation of various metabolic pathways. However, its biological relevance remains uncertain due to methodological difficulties in determining high-impact succinylation sites. In the present study, using stable isotope labeling and data-independent acquisition mass spectrometry, we quantified lysine succinylation stoichiometries in mouse livers. Despite the low overall stoichiometry of lysine succinylation, several high stoichiometry sites were identified, especially upon deletion of the desuccinylase SIRT5. In particular, multiple high stoichiometry lysine sites identified in argininosuccinate synthase (ASS1), a key enzyme in urea cycle, are regulated by SIRT5. Mutation of the high stoichiometry lysine in ASS1 to succinyl-mimetic glutamic acid significantly decreased its enzymatic activity. Metabolomics profiling confirms that SIRT5 deficiency decreases urea cycle activity in liver. Importantly, SIRT5 deficiency compromises ammonia tolerance and reduces locomotor and exploratory activity in male mice upon high-ammonium diet feeding. Therefore, lysine succinylation is functionally important in ammonia metabolism.

## Introduction

Lysine succinylation is a type of protein post-translational modification (PTM)^1,2^. The acidic succinyl group alters the structure and function of modified proteins by increasing the size of modified lysine residues and converting the positive charge on the ε-amino group to a negative charge. Using mass spectrometry (MS), often in conjunction with immunoaffinity enrichment, thousands of succinylated lysine sites have been identified across prokaryotes and eukaryotes. These succinylated proteins are widely involved in metabolic pathways, including the tricarboxylic acid (TCA) cycle, the urea cycle, fatty acid oxidation, and ketone body synthesis pathways^3-6^. Thus, succinylation plays multiple potentially important roles in the regulation of metabolism and is implicated in the pathogenesis of multiple diseases^7,8^.

Succinyl-CoA, a metabolic intermediate in the TCA cycle, is the endogenous donor of succinyl groups. Several enzymes were recently reported to possess a “moonlighting” succinyltransferase activity, and regulate lysine succinylation towards specific substrates^9,10^. By and large, however, due to the intrinsic reactivity of succinyl-CoA, lysine succinylation is non-enzymatically formed^11^. Through protein succinylation, chronic exposure to succinyl-CoA may cause cumulative cellular damage that contributes to aging and diseases. Succinylation is therefore considered as a form of “carbon stress” that threatens proteome integrity^12^. To counter this process and restore protein function, succinylation is selectively targeted for removal by sirtuin 5 (SIRT5)^1,13^, a member of the nicotinamide adenine dinucleotide (NAD^+^)-dependent deacylase sirtuin family. The desuccinylase activity of SIRT5 may therefore underlie some of the health benefits of NAD^+^ and calorie restriction.

So far, studies have assessed relative changes of lysine succinylation sites after antibody enrichment using MS-based proteomics^3-6^. Although determining relative succinylation changes is important, this does not reveal site occupancy, or stoichiometry, of individual modification sites, which is critical to understand the functional consequences of succinylation (“succinylation stoichiometry” at a particular lysine site is defined as the fraction of the total amount of a protein succinylated on the site to the total amount of that protein). Among acylation modifications, lysine acetylation is the simplest and best characterized type. It has been reported in both yeast and mammals that most acetylation lysine sites are of low stoichiometries (<1%) ^14-18^. These observations question the functional importance of most acetylation sites and cast doubts on the biological relevance of lysine acylations in general. Despite the potentially dramatic structure alteration upon succinylation and the increasing number of identified succinylated proteins, due to a poor understanding of its occupancies, the biological relevance of lysine succinylation remains largely uncertain. To solve this problem, in the present study, we quantified lysine succinylation stoichiometries in mouse liver to determine the functional importance of succinylated proteins.

## Results

### Succinylome profiling using GluC for protein digestion

The quantification of succinylation stoichiometry depends on the accurate determination of endogenously succinylated proteins and non-succinylated proteins. To distinguish succinylated and unmodified lysine residues, we employed a heavy isotope labeling method by per-succinylation of proteins with deuterated succinic anhydride-*d*_4_, so that peptide isoforms generated *in vivo* (endogenous, light isoform) and *in vitro* (exogenous, heavy isoform) can be determined by liquid chromatography–mass spectrometry (LC-MS/MS) and data analysis (as detailed later in **Fig. 2**). In previous studies on lysine succinylation^3-6^, trypsin has been used for protein digestion in sample preparation for LC-MS. However, trypsin, which cleaves peptide chains mainly at the carboxyl side (C-terminal) of lysine or arginine residues, cannot cut efficiently at the C-terminal side of acylated lysine residues. Thus, tryptic digestions of per-succinylated proteins typically generate peptides that are very large and often contain many succinylated lysine residues, complicating analysis. GluC, however, is an endoproteinase specific for C-terminal cleavage at glutamic and aspartic acid residues, and the cleavage pattern or efficiency of GluC should not be impacted by the presence of lysine succinylation. Therefore, GluC was used for the digestion of per-succinylated proteins in this study.

To generate a reference lysine succinylome for the quantification of succinylation stoichiometry in mouse liver, we developed a workflow to enrich succinylated peptides prepared with GluC digestion for succinylation identification by MS (**Fig. 1a**). Mouse liver protein samples were isolated from four wildtype (WT) and four *Sirt5*^*-/-*^ mice. Equal amounts of protein from each sample of WT and knockout (KO) mice were digested with GluC. Succinylated peptides were enriched by immunoaffinity and analyzed in duplicate by LC-MS/MS on a TripleTOF 6600 mass spectrometer, and data were searched against the *Mus musculus* protein database using ProteinPilot. We identified 1104 lysine succinylation sites across 298 proteins with a false discovery rate (FDR) of 1% (**Fig. 1b**, and **Supplementary Data 1**). Of the 1104 sites identified, 505 (46%) were identified only in the KO, 175 (16%) only in the WT, and 424 (38%) in both (**Fig. S1**). These percentages are comparable to our previously generated mouse liver mitochondrial succinylome using tryptic digestion^4^, and confirms the importance of SIRT5 in regulating the overall lysine succinylation. Interestingly, when we accessed the individual succinylated peptides and sites identified in this and previous studies, we observed many novel succinylation sites and relatively low overlap between the succinylomes generated by tryptic *versus* GluC digestion (**Fig. 1b** and **Fig. S2a**). Approximately 30% of succinylation sites identified in mitochondrial proteins through tryptic digestion in our previous study^4^ were identified using GluC as protease in this study. Similarly, only 20% of succinylation sites identified by Park et al.^3^ in mouse liver after tryptic digestion were identified by using GluC here. Importantly, 673 succinylation sites (61%) identified through GluC digestion in this study are reported for the first time, and have not been identified by using tryptic digestion before (**Fig. 1b**). This observation suggests the limitation of trypsin-based proteomics study in PTM discovery, and shows a clear benefit of using a different proteolytic enzyme to increase overall succinylome coverage.

**Fig. 1.**
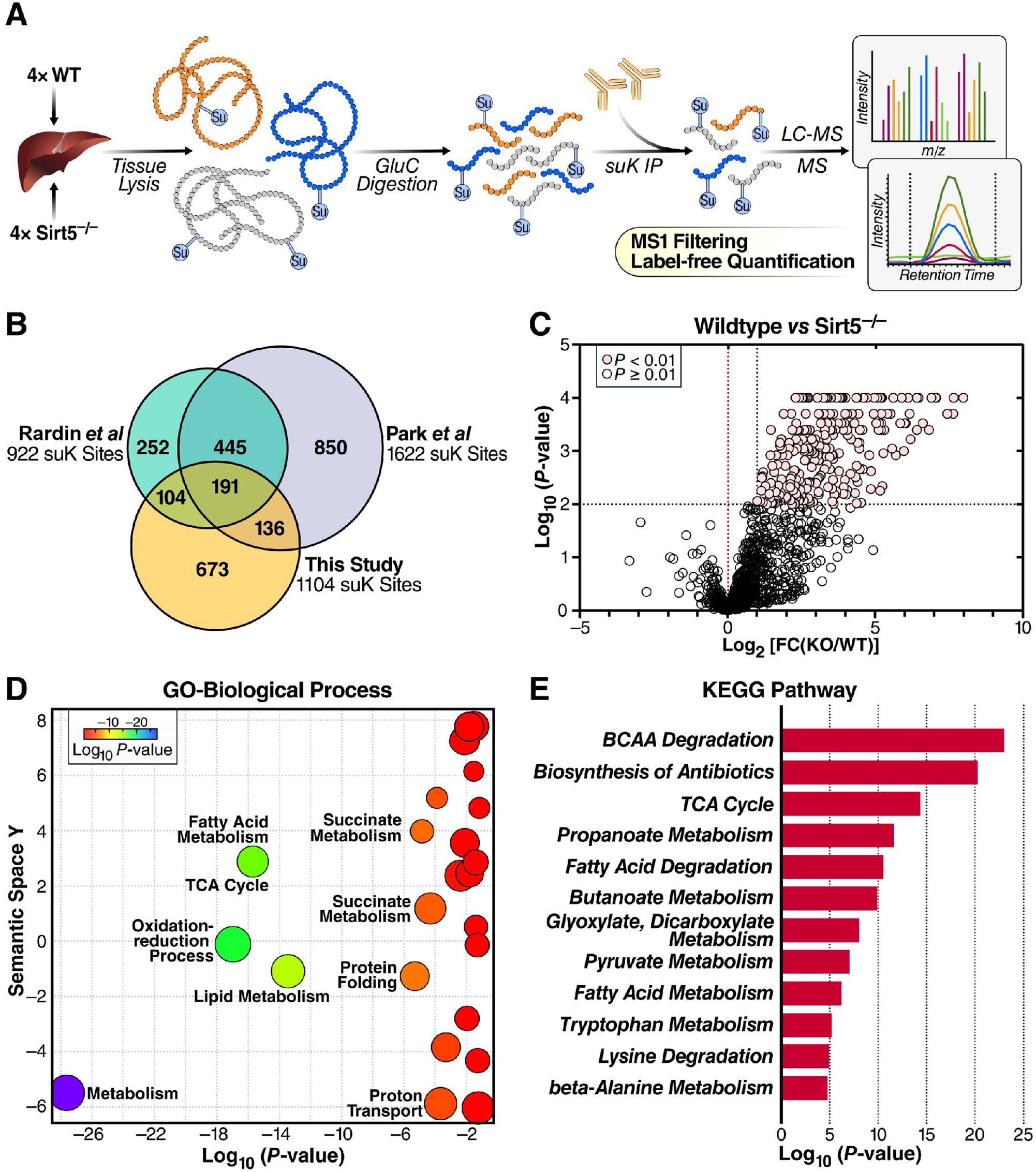
Enrichment and identification of liver lysine succinylome. **(a)** Workflow of enrichment and label-free quantitation of lysine succinylome in WT and *Sirt5*^*-/-*^ mouse livers. **(b)** Venn diagrams of succinylated lysine sites identified in this study (WT and *Sirt5*^*-/-*^ mouse livers) and previous studies (Rardin et al., *Cell Metab*. 2013; Park et al., *Mol Cell*. 2013)^3,4^. **(c)** Volcano plot showing the differential regulation of the identified succinylated sites. **(d)** GO analysis of succinylated proteins that were targeted by SIRT5 (> twofold increase and p < 0.01) using DAVID and REVIGO. **(e)** KEGG pathway analysis of succinylated proteins that were targeted by SIRT5 (> twofold increase and p < 0.01) using ToppFun. Top 12 most-enriched pathways are shown here.

To investigate what proteins in this GluC digestion-based immunoaffinity-enriched succinylome are regulated by the deacylase SIRT5, we quantified the fold-changes of lysine succinylation upon SIRT5 KO using Skyline. The results showed that 225 (23%) lysine sites on 104 (37%) succinylated proteins showed greater than twofold change of succinylation in the SIRT5 KO mice, compared to WT. As expected, the majority of these SIRT5-targeted sites had higher succinylation levels in the SIRT5 KO group (**Fig. 1c**, and **Supplementary Data 1**). Functional analyses suggest that SIRT5-targeted proteins were involved in biological processes, including oxidation-reduction process, fatty acid metabolism, lipid metabolism, and TCA cycle (**Fig. 1d**). KEGG pathway analysis confirmed that the SIRT5-targeted proteins are enriched in major metabolic pathways, including branched chain amino acid (BCAA) degradation, biosynthesis of antibiotics, TCA cycle, propanoate metabolism, and fatty acid degradation, among many others (**Fig. 1e**). Next, considering the small overlap of tryptic and GluC-based succinylomes, we wanted to know which pathways are commonly identified by both GluC- and trypsin-based proteomes, and which pathways are specifically enriched in the present GluC-based proteome. Comparative analysis of SIRT5-targeted (fold-change > 2, and p < 0.01) succinylated proteins identified in this and our previous study^4^ showed that most of the enriched KEGG pathways generated in the two studies are the same, with only slight differences in the ranks of enrichment P values (**Fig. S2b**). This observation is not surprising because the pathways enriched appear rather robustly regulated, and usually several enzymes of a given pathway are strongly regulated. The above results suggest that succinylated proteins enriched in major metabolic pathways are widely targeted by SIRT5, which necessitates the quantification of succinylation stoichiometry to further determine their functional significance.

### Quantification of lysine succinylation stoichiometry in mouse liver

To quantify lysine succinylation stoichiometry, we developed a novel method by modifying the stable isotope–labeling method from Baeza et al.^19^ and Meyer et al.^20^, followed by SWATH acquisition that collects both precursor and multiple fragment ion abundances^20,21^. Briefly, protein lysates were extracted from whole-liver tissues derived from four WT and four *Sirt5*^*-/-*^ mice (**Fig. 2**). Second, protein lysates were incubated three times with succinic anhydride-*d*_4_ to succinylate unmodified lysine residues. Third, samples were digested with endoproteinase GluC, and the proteolytic peptides were fractionated by basic reversed-phase chromatography. Finally, peptides were analyzed by LC-MS in SWATH acquisition mode. Only ions containing the modified lysine of interest were used for quantification, as those can be used to differentiate between the light and heavy succinylated peptide forms. Stoichiometry measurements were computed from light (L) and heavy (H) peak areas as the percentage of L/(L + H).

**Fig. 2.**
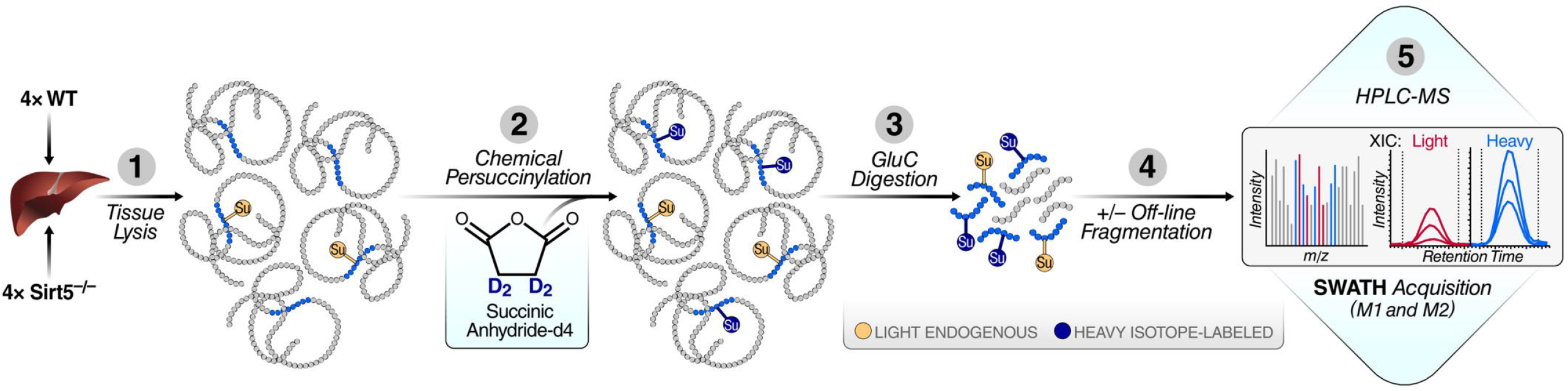
Succinylation stoichiometry workflow. Proteins in clarified tissue lysate are incubated with succinic anhydride-d_4_ to succinylate unmodified lysine residues. Next, acetylated proteins are digested with endoproteinase Glu-C, followed by HPLC fractionation of the proteolytic peptides using basic-pH reversed-phase chromatography. Finally, peptides are analyzed by LC-MS using DIA with variable precursor window widths. Pink circles indicate light succinyl groups, and blue circles indicate heavy succinyl groups.

By using this method, we identified 822 succinylated lysine sites in mouse liver, and quantified their succinylation stoichiometries in both WT and SIRT5 KO samples (**Supplementary Data 2**). The results showed a wide range of stoichiometries across succinylated sites from nearly 0 to over 90%; average stoichiometries of these sites are 5.25% in WT, and 6.11% in KO livers (**Fig. 3a**). However, many of these identified succinylated lysines may have been only chemically modified with heavy labeled succinyl group, and do not genuinely exist *in vivo*. To remove these non-endogenously succinylated sites, we turned to the reference succinylome generated through immunoaffinity enrichment and GluC digestion described earlier (**Fig. 1**). By comparing with the immunoaffinity-enriched succinylome, we narrowed the succinylated sites in the stoichiometry dataset down to 238 sites (on 103 proteins) of high confidence that were believed to exist endogenously (**Fig. 3a**). The average stoichiometry of these high-confidence sites was 3.57% (median: 1.58%) in WT and 4.05% (median: 1.68%) in KO livers (**Fig. 3a**). The majority of these sites have a succinylation stoichiometry of 0.1–5%, and only 41 sites have a succinylation stoichiometry >5% in at least one of the two genotypes (**Fig. 3b**). Although most lysine sites have comparable levels of succinylation stoichiometries in WT and SIRT5 KO livers, some sites, including ASSY (K112 and K121), ECI2 (K301), PYC (K35), and HCDH (K192), showed a remarkably higher succinylation stoichiometry in SIRT5 KO liver than in WT liver (high ΔStoichiometry) (**Fig. 3c** and **3d**). The succinylation on these sites is thus more likely to be regulated by SIRT5 and biologically relevant.

**Fig. 3.**
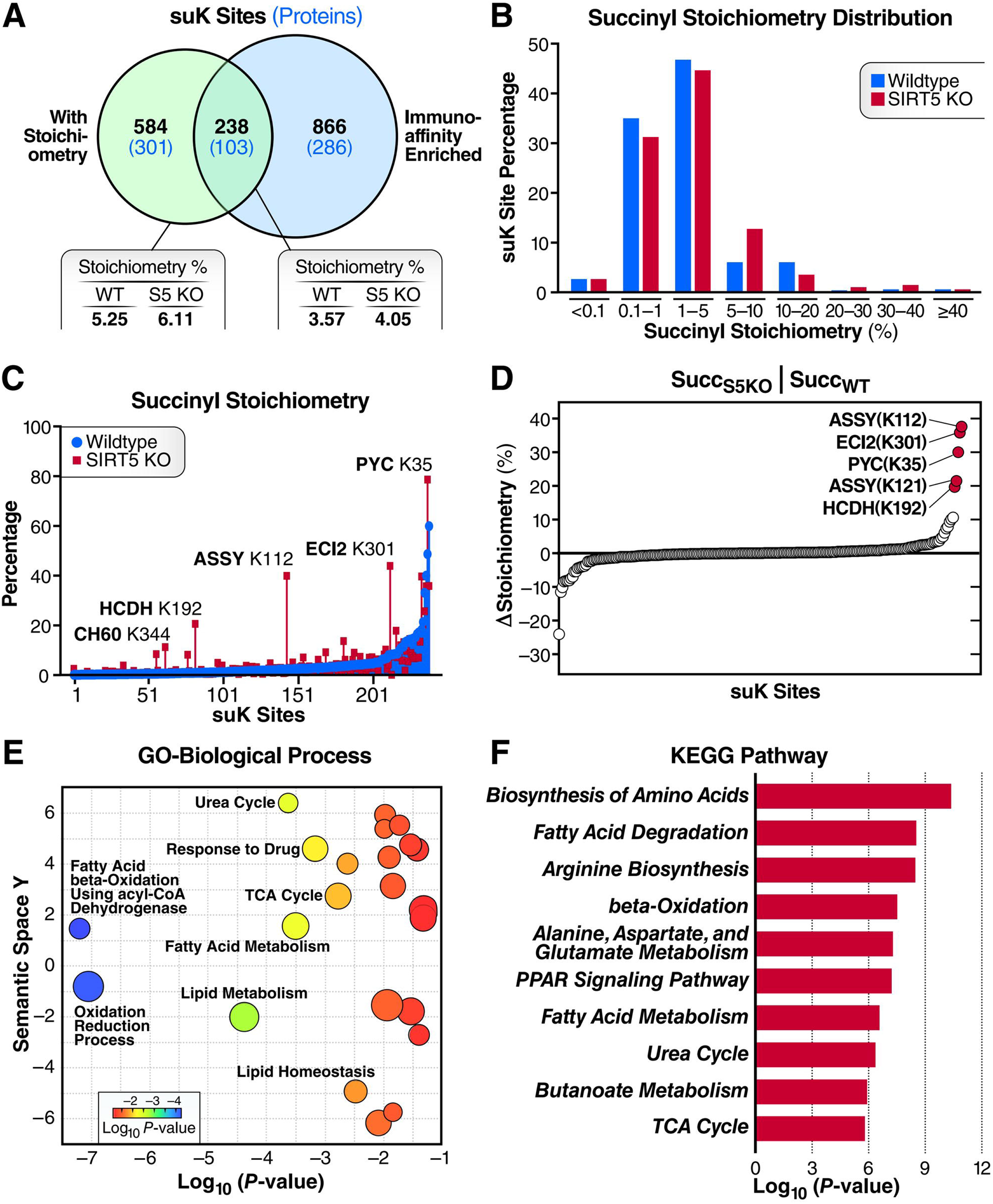
Quantification of lysine succinylation stoichiometry in mouse liver. **(a)** Venn diagram showing succinylated lysine sites and proteins (in parenthesis) identified in the stoichiometry MS dataset and the succinylation immunoaffinity-enrichment MS data. Average stoichiometries are indicated in boxes. **(b)** Distributions of SuK stoichiometries between WT and *Sirt5*^*-/-*^ mouse livers. **(c)** Stoichiometries of SuK sites in WT and *Sirt5*^*-/-*^ mouse livers. **(d)** Δ Stoichiometry (*Sirt5*^*-/-*^ (S5KO) - WT) of confirmed SuK sites. **(e)** GO analysis of proteins with succinylation stoichiometries higher than 5% using DAVID and REVIGO. **(f)** KEGG pathway analysis of proteins with succinylation stoichiometries higher than 5% using ToppFun. Top 12 most enriched pathways are shown here.

To identify which biological processes or pathways could be regulated through lysine succinylation, we performed functional analyses of high stoichiometry (>5%) lysine sites. Gene ontology analysis showed that high stoichiometry sites are mostly enriched in fatty acid beta oxidation, oxidation-reduction process, lipid metabolism, and the urea cycle (**Fig. 3e**). KEGG pathway analysis confirmed the enrichment of most of these biological processes (**Fig. 3f**). Amino acid metabolic pathways (in particular, arginine biosynthesis pathway; alanine, aspartate and glutamine metabolism pathway; and urea cycle pathway) are among the pathways that high stoichiometry proteins were most significantly enriched in (**Fig. 3f**), implying that lysine succinylation plays an important role in regulating nitrogen metabolism in the liver.

### Mutation at a high stoichiometry succinylation site of ASS1 affects its thermal stability and enzymatic activity

Among succinylated proteins identified in the proteomics dataset, argininosuccinate synthase (ASSY, also known as ASS1) had lysine sites with the highest SIRT5-regulated succinylation stoichiometries (ΔStoichiometry). The succinylation stoichiometries of K112 and K121 are remarkably higher in SIRT5 KO mouse liver (39.9% and 39.6%, respectively) than in WT controls (2.3% and 18.1%, respectively) (**Fig. 4a**), we next sought to investigate if the succinylation of these lysine residues is functionally important.

**Fig. 4.**
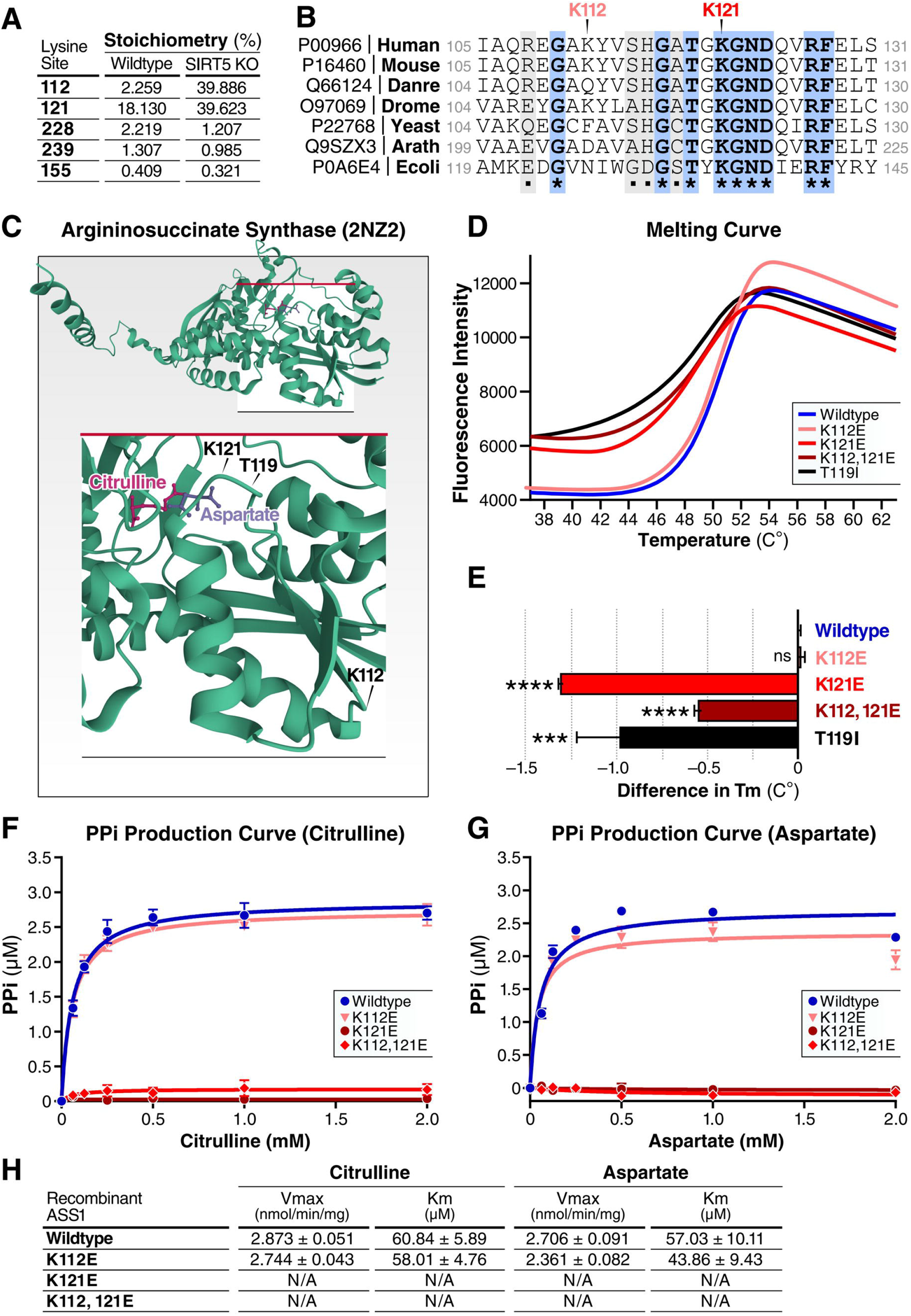
Mutation at a high-stoichiometry succinylation site of ASS1 affects its thermal stability and enzymatic activity. **(a)** Succinylation stoichiometries of lysine sites on ASS1 in WT and *Sirt5*^*-/-*^ mouse livers that were confirmed by immuno-affinity enrichment proteomics. **(b)** Alignment of protein sequences of argininosuccinate synthases across species using UniProt. Completely conserved (*) and partially conserved (.) sites are highlighted. **(c)** Cartoon of the crystal structure of human ASS1 (PDB entry 2NZ2) bound with aspartate and citrulline. **(d)** Thermostability of WT and mutant recombinant human ASS1 proteins examined by differential scanning fluorimetry. **(e)** Changes in melting temperature (Tm) values compared with that of the WT ASS for each ASS1 mutant. Error bars depict s.e.m. from four measures. **(f-h)** Steady-state kinetic analysis of WT and mutant ASS1 enzymatic activity as measured by the production of pyrophosphate from increasing concentrations of aspartate **(f)** and citrulline **(g)**. Graph is representative of two independent experiments, n = 4 measurements/sample, mean ± s.d.. Values are shown for average Km and Vmax ± s.e.m. **(h)**.

Argininosuccinate synthase is an evolutionarily conserved enzyme catalyzing the synthesis of argininosuccinate (AS) from citrulline and aspartic acid. Alignment of the amino acid sequences of argininosuccinate synthase proteins in different organisms showed that K121 is highly conserved across the evolutionary spectrum ranging from prokaryotes (*E. coli*) to mammals and plants (*Arabidopsis thaliana*), and K112 is less conserved (**Fig. 4b**). Structurally, K121 is located at the binding pocket for citrulline and aspartic acid, whereas K112 is located within a non-structured segment distant from the binding pocket (**Fig. 4c**), which implicates a potentially more critical role for K121 in ASS1 function.

Next, we constructed recombinant human ASS1 proteins with a lysine (K) to glutamic acid (E) mutation at either K112 or K121 or at both sites to mimic succinylation on these sites. Because T119 is an aspartic acid binding site, and T119I mutation was identified in patients with Type I citrullinemia^22^, we also generated a threonine (T) to isoleucine (I) mutant on T119 as a pathogenic construct control. We purified these mutant and WT recombinant ASS1 proteins and examined their thermal stability by measuring their melting curves using differential scanning fluorimetry. The results showed that, as expected, the T119I mutation caused a left shift of the melting curve (**Fig. 4d**), indicating that the pathogenic T119I mutation reduces the temperature required for ASS1 denaturation. Similarly, we observed that both the K121E single mutation and the K112,121E double mutation caused a left shift of the melting curve of ASS1 (**Fig. 4d**), indicating that these mutations reduce the denaturation temperature of ASS1. However, the K112E single mutation did not overtly change the melting curve (**Fig. 4d**). Quantification of the changes in melting temperature (Tm) values, compared with that of the WT ASS1, for each ASS1 mutant confirmed that the T119I, K121E, and K112,121E mutations reduce the Tm values, whereas the K112E mutation did not significantly change ASS1 Tm value (**Fig. 4e**). These observations suggest that the succinylation-mimic K-to-E mutation on K121, but not K112, decreases ASS1 thermal stability.

To further determine the importance of K112 and K121 on ASS1 enzymatic activity, we measured the kinetics of the recombinant ASS1 proteins for its substrates, aspartic acid and citrulline, respectively. Steady-state kinetic analysis of the WT and K-to-E mutant ASS1 was performed by measuring the production of pyrophosphate (PPi) from increasing concentrations of the substrate. The results showed that the K112E mutation did not significantly change the Km values of ASS1 for both citrulline and aspartic acid, whereas the K121E single mutation or the K112,121E double mutation almost completely removed the activity of ASS1 (**Fig. 4f-h**). Consistently, K112E mutation only slightly reduced the Vmax value of ASS1 for both substrates, whereas the K121E and K112,121E mutants did not have a measurable Vmax value (**Fig. 4f-h**). These results suggest that K121, but not K112, is critical for ASS1 activity, and support the notion that succinylation of K121 in ASS1 with a high stoichiometry disrupts ASS1 activity and may compromise urea cycle function at large.

### SIRT5 deficiency affects metabolites in urea cycle in mouse livers

To evaluate the metabolic consequence of succinylation, we performed metabolomic profiling in mouse livers. Metabolites extracted from eight WT and eight *Sirt5*^*-/-*^ mouse livers were analyzed by LC/MS. A total of 367 metabolites were detected and quantified in both groups (**Supplementary Data 3**). Principal component analysis showed that WT and SIRT5 KO groups were largely separated, suggesting a functional impact of SIRT5 on the overall hepatic metabolome (**Fig. 5a**). Among the 367 quantified metabolites, eight were significantly upregulated upon SIRT5 KO (p < 0.05), and 49 were downregulated by SIRT5 KO. Notably, ornithine, a urea cycle intermediate metabolite, was among the most downregulated metabolites in SIRT5 KO mouse livers, compared with WT controls (**Fig. 5b**). Functional enrichment analyses showed that the metabolites upregulated upon SIRT5 KO were enriched in pathways, such as the glycerol phosphate shuttle, *de novo* triacylglycerol biosynthesis, and cardiolipin biosynthesis, among others (**Fig. 5c**), and metabolites downregulated by SIRT5 KO were mostly enriched in arginine and proline metabolism, taurine and hypotaurine metabolism, and the urea cycle (**Fig. 5d**). This observation is consistent with the stoichiometric succinylome results, which showed that arginine biosynthesis, alanine, aspartate and glutamine metabolism, and urea cycle are among the pathways that high stoichiometry proteins were most significantly enriched in (**Fig. 3f**). Specifically, metabolomics results showed that argininosuccinate (the product of ASS1) and its downstream metabolites, arginine and ornithine, were all significantly lower in SIRT5 KO mouse livers than WT controls (**Fig. 5e**). Although not statistically significant, other metabolites involved in urea cycle, such as aspartic acid and fumaric acid, also trended lower in SIRT5 KO samples (**Fig. 5e**). These results confirmed a regulatory role of SIRT5 in urea cycle at the metabolite level, and support the functional importance of succinylation of urea cycle enzymes, including ASS1.

**Fig. 5.**
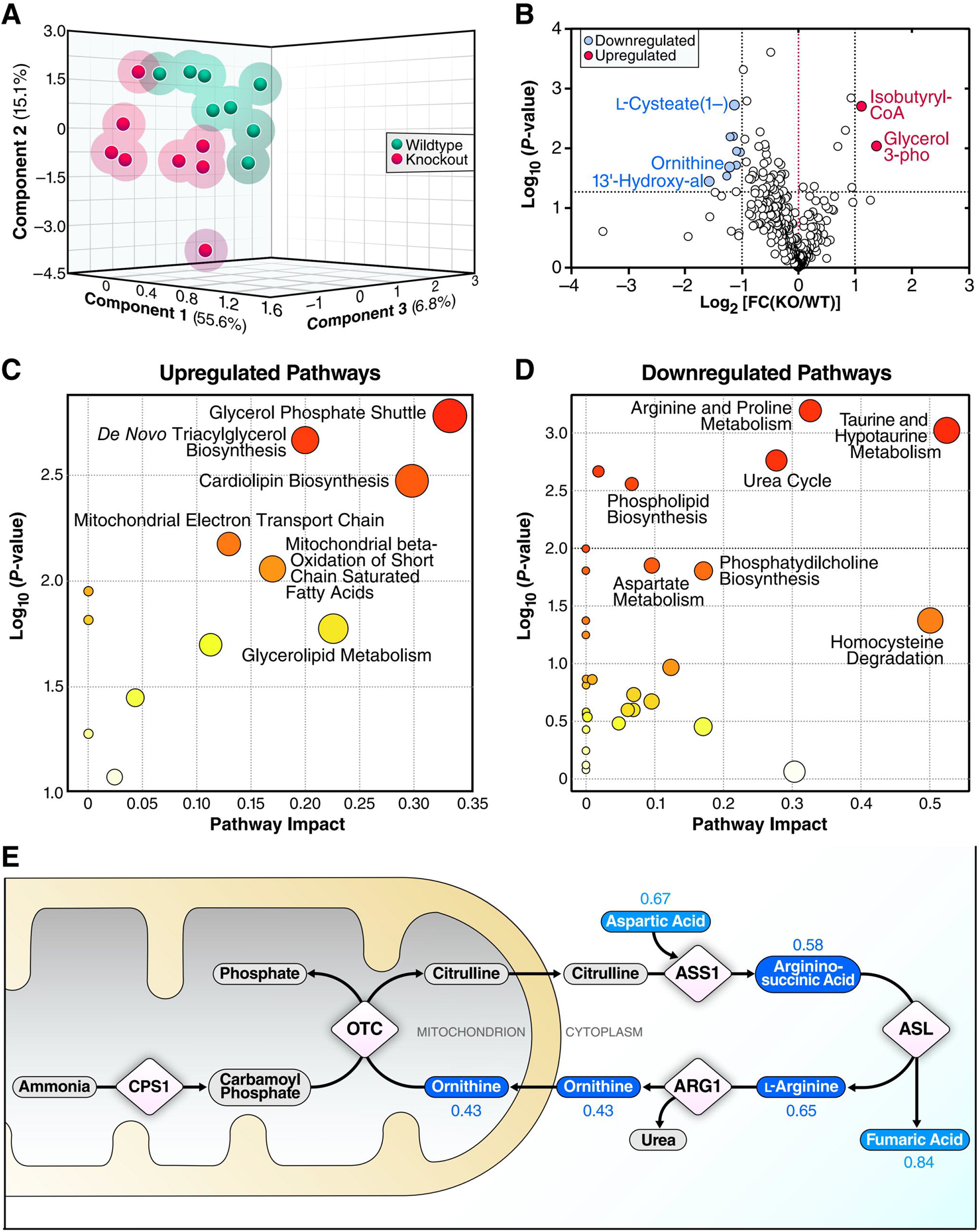
SIRT5 deficiency affects metabolites in urea cycle in mouse livers. **(a)** Component analysis of hepatic metabolome showing the relative distance between WT and *Sirt5*^*-/-*^ samples, n = 8. **(b)** Volcano plot showing the differential abundance of the identified metabolites without functional SIRT5. **(c)** SMPDB pathway analysis of upregulated metabolites (p < 0.05) using MetaboAnalyst. **(d)** SMPDB pathway analysis of downregulated metabolites (p < 0.05) using MetaboAnalyst. **(e)** Schematic diagram of urea cycle showing downregulated (light blue capsules: p ≥ 0.05; dark blue capsules: p < 0.05), and unidentified (gray capsules) metabolites. KO:WT ratios are indicated above the metabolites in the corresponding colors.

### SIRT5 deficiency reduces ammonia detoxification and locomotor activity upon high-ammonium diet feeding in male mice

To further examine SIRT5’s effect on ammonia detoxification, we measured blood ammonia levels in 21-month-old SIRT5 KO homozygous (*Sirt5*^−/-^), heterozygous (*Sirt5*^+/-^), and WT mice. SIRT5 KO and *Sirt5*^+/-^ mouse plasma ammonia trended higher than WT littermates on standard chow diet (**Fig. 6a**). Next, we challenged the ammonia detoxifying function in 3-month-old SIRT5 KO and WT male mice by putting them on a high-ammonium diet (HAD) for 4 weeks. The HAD, which has been used to induce hyperammonemia in mice and rats, was formulated by adding 20% ammonium acetate into a chow diet (control diet)^23,24^. During the 4 weeks of special diet feeding, food intake was monitored every week. The results showed that food intake in HAD-fed group was about 53–85% of that in the control group (**Fig. S3a**). Considering that the calories in HAD were 20% lower than that in the same amount of control diet, the calories that HAD-fed mice ingested was only about 42–68% of what control diet-fed mice ingested on average. Therefore, while the control diet-fed mice were gradually gaining weight, the body weights of their HAD-fed litter mates were continually decreasing (**Fig. S3b**). After 4 weeks of the special diet feeding, plasma ammonia levels were measured. The results showed that plasma ammonia levels were once again comparable in these young SIRT5 KO and WT mice fed on a control diet (**Fig. 6b**). However, WT mice showed the trend of a lower level of plasma ammonia upon HAD feeding (**Fig. 6b**), possibly due to a compensatory increase in ammonia detoxification protein expression. On the contrary, SIRT5 KO mice had a higher plasma ammonia level after HAD feeding (**Fig. 6b**). These results suggest that 4 weeks of HAD-feeding was well tolerated in WT mice, whereas SIRT5 deficiency reduced ammonia tolerance upon HAD feeding.

**Fig. 6.**
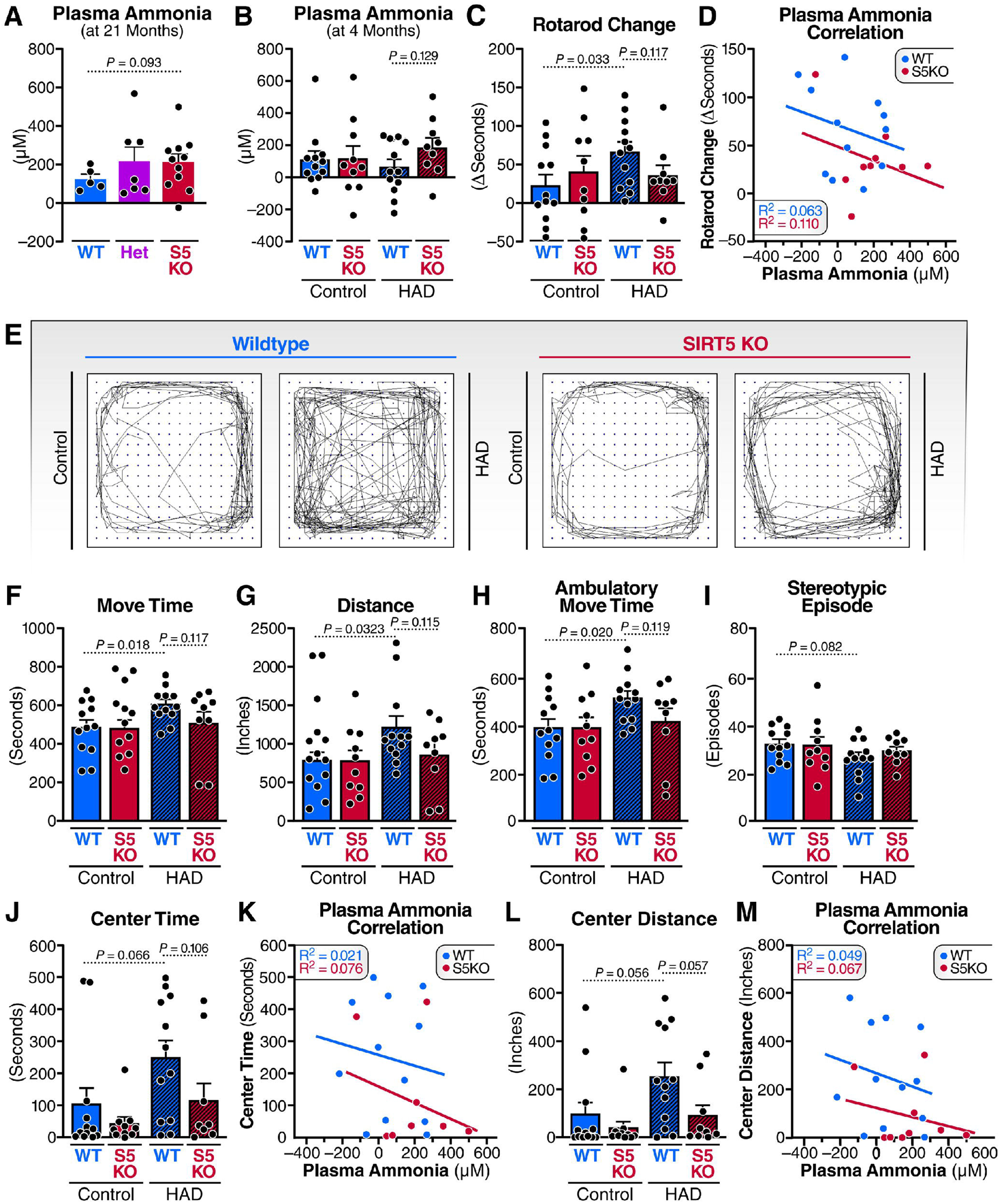
SIRT5 deficiency reduces ammonia detoxification and locomotor activity upon high-ammonium-diet feeding in male mice. **(a)** Ammonia levels measured in the plasma from 21-month-old WT (n = 5), *Sirt5*^*+/-*^ (Het, n = 7) and *Sirt5*^*-/-*^ (S5KO, n = 11) mice after 24 h of fasting. Error bars indicate mean ± s.e.m.. P value was generated with unpaired student t test. **(b)** Ammonia levels measured in the plasma from 4-month-old WT and *Sirt5*^*-/-*^ (S5KO) male mice after being fed on a chow diet (control) or high-ammonium diet (HAD) for 4 weeks without fasting. n = 9–12. Error bars indicate mean ± s.e.m.. **(c)** Per-mouse change in rotarod assay performance (time on rotarod on Day 28 – time on rotarod on Day 0). n = 9–12. Error bars indicate mean ± s.e.m.. P values were generated with unpaired student t test. **(d)** Correlation between rotarod performance improvement and plasma ammonia levels in WT and S5KO male mice fed on HAD was tested via linear regression. **(e-m)** Open-field testing of WT and S5KO male mice was performed after 4 weeks of control diet or HAD feeding. Representative tracks of paths are shown in (**e**). Movement time (**f**), total distance (**g**), ambulatory movement time (**h**), stereotypic episode (**i**), and center time (**j**) and center distance (**l**) were calculated. Correlation between center time and plasma ammonia level (**k**), and correlation between center distance and plasma ammonia level (**m**), in WT and S5KO male mice fed on HAD, was tested via linear regression. n = 9-12. Error bars indicate mean ± s.e.m.. P values were generated with unpaired student t test.

Ammonia toxicity is a prominent cause of hepatic encephalopathy^25^. In consideration of the link between hyperammonemia and impairment in motor and cognitive function^26^, we examined locomotor coordination and activity in SIRT5 KO and WT mice. Rotarod test results showed that average performance improved over the 4 weeks of control diet or HAD feeding, and the improvement in rotarod performance was negatively correlated with plasma ammonia level, in both WT and SIRT5 KO mice (**Fig. S3e**). Consistent with the decreased plasma ammonia level in WT mice upon HAD feeding, HAD-feeding increased the improvement in rotarod performance in WT mice (**Fig. 6c-d**). By contrast, the improvement in rotarod performance was not increased after HAD feeding in SIRT5 KO mice (**Fig. 6c**), which is in line with their increased blood ammonia level (**Fig. 6d**). Grip strength test results showed that the limb muscle strength of WT and SIRT5 KO mice were comparable (**Fig. S3c-d**), suggesting that the lower level of improvement in rotarod performance in SIRT5 KO mice was not due to lower muscle strength in SIRT5 KO mice, and that SIRT5 deficiency reduces improvement in coordination and balance in male mice fed on HAD.

Next, we performed an open-field test to assess locomotor activity and exploratory behavior of the mice. The results showed that move time, move distance, and the time of exploring in the center of the arena were all comparable in SIRT5 KO and WT mice on a control diet (**Fig. 6e-m**), suggesting that SIRT5 deficiency does not affect locomotor activity and exploration under a baseline condition. By contrast, the results showed that HAD-feeding increased the move time and distance in WT mice, but failed to increase movement in SIRT5 KO mice (**Fig. 6e-g**). Results also showed that the HAD-induced movement could be attributed to the increase in ambulatory movement, but not stereotypic movement (repetitive behavior) (**Fig. 6h-i**). These results suggest that SIRT5 deficiency reduced the increase in locomotor activity upon HAD feeding. Furthermore, the center time and distance were increased by HAD-feeding in WT mice, but not in SIRT5 KO mice (**Figs. 6j and 6l**), suggesting that SIRT5 KO mice had a higher level of anxiety and were less exploratory than WT mice on a HAD. Regression analysis confirmed the negative correlation between plasma ammonia level with center time and distance (**Figs. 6k, 6m, and S3f-g**). These results provide *in-vivo* evidence supporting an important role of SIRT5 in ammonia detoxification and locomotor activity.

## Discussion

In the present study, we profiled the liver succinyl proteome using GluC digestion coupled with LC-MS/MS using SWATH acquisition, and quantified the site-specific succinylation stoichiometry in mouse liver using an isotope labeled chemical succinylation approach. Since the functional analysis of high-stoichiometry succinylated sites pointed towards an enriched urea cycle pathway, we further studied the importance of high-stoichiometry lysines and found that succinylation mimicking K-to-E mutation in one of the most affected high stoichiometry urea cycle enzymes, ASS1, disrupts its enzymatic activity. Subsequent metabolomic profiling and behavioral testing upon HAD-feeding confirmed the decline of urea cycle function and ammonia detoxification in SIRT5 KO mouse liver. This is, to our knowledge, the first time succinylation stoichiometry was quantified at a proteomic scale in a mammalian tissue, and the results highlight the urea cycle as a major metabolic process in liver being functionally modulated by lysine succinylation.

In the past decade, since the discovery of lysine succinylation and the desuccinylase activity of SIRT5^1,2^, thousands of succinylated proteins have been identified in different organisms. Lysine succinylation has thus been implicated in the regulation of a wide range of biological processes and metabolic pathways, including the TCA cycle, the urea cycle, fatty acid β-oxidation, and ketone body synthesis^3-6^ to just name a few. The quantification of succinylation levels in these studies was based on the relative fold-change under different conditions, and these differentially succinylated peptides are biologically relevant only if the succinylation occupancies on one or more specific lysine sites are high enough to affect the function of the modified proteins. A large increase in fold-change would thus not create a phenotypic difference if the absolute stoichiometry of succinylation is very low. Therefore, quantification of succinylation stoichiometries is required to inform the functional importance of succinylated proteins.

Kinase signaling pathways regulate biological processes in a switch-like “on-and-off” manner; in many cases, phosphorylation shifts between near-zero and nearly full occupancy states when modified within a given compartment. Proteome-scale studies confirmed this notion by showing that about one out of 10 phosphorylation sites have a very high (>90%) phosphorylation occupancy in exponentially growing yeast^27^, and that nearly half of all phosphorylation sites have an occupancy of 70% or higher in HeLa cells during mitosis^28^. By contrast, acetylation, the chemically simplest acylation, appears to have a generally much lower stoichiometry. By using heavy isotope labeled acetic anhydride to chemically label unmodified lysine, Baeza et al. quantitatively calculated the average global acetylation stoichiometry was about 7% in *E. coli*, and that only 4% of the quantified acetylated lysine sites had a stoichiometry greater than 20%^19^. By using a similar strategy, Zhou et al. reported that the average lysine acetylation stoichiometry is about 4.1%, and that only 4.8% of acetylated sites had stoichiometries over 20% in HeLa cells^18^. Meanwhile, by using partial chemical acetylation and stable isotope labeling by amino acids (SILAC)-based quantification, Weinert et al. reported that 95% of acetylation sites had a stoichiometry <1% in exponentially growing yeast^14^. With the same approach, they also estimated that the median stoichiometry of acetylation was only ∽0.05% in mouse liver^15^, and ∽0.02% in HeLa cells^16^. These studies suggest that the majority of acetylated lysine sites are of very low acetylation stoichiometries, and not biologically important. As for lysine succinylation, Park et al. adopted a mathematical approach and calculated succinylation stoichiometries based on the SILAC ratios of modified peptides, unmodified peptides and protein ratios in mouse embryonic fibroblasts^3^. This mathematical approach is very useful, but indirect estimation of succinylation stoichiometry may not be accurate or robust due to the amplification of errors over the multiple calculation steps.

In the present study, we developed a novel workflow by performing heavy isotope–labeled chemical per-succinylation of unmodified lysine residues followed by quantitative MS to measure succinylation stoichiometry in mouse livers. Through this approach, we quantified succinylation stoichiometries on 238 lysine sites with high confidence in 103 proteins (high-confidence succinylation sites are the ones confirmed with GluC digestion-based immunoaffinity-enrichment succinyl proteome). We report that the stoichiometries of succinylation on these sites are 3.57% on average, and the median stoichiometry is 1.58%, in WT mouse livers. In SIRT5 KO mouse livers, the stoichiometries on these sites are 4.05% on average, and the median stoichiometry is 1.68%. Our study therefore suggests that, similar to lysine acetylation, the occupancy of succinylation is very low in most modified substrates, and only a small group of high-stoichiometry proteins appear to be functionally regulated by lysine succinylation.

Through functional enrichment analysis of high stoichiometry (>5%) sites, we find that the urea cycle pathway was among the most enriched metabolic pathways. In particular, the high succinylation stoichiometry lysine sites in ASS1 were remarkably higher in SIRT5 KO livers. We further determined that, although both sites have high succinylation stoichiometries upon SIRT5 KO, the succinylation of K121, but not K112, is critical for a normal ASS1 function, because a succinylation-mimicking K-to-E mutation of K121, but not K112, reduces ASS1 thermal stability, and almost completely deprives ASS1 activity. Thus, ASS1 appears to be an important succinylated substrate of SIRT5 involved in the regulation of the urea cycle in mouse liver. Consistently, hepatic metabolomics profiling showed reduced levels of urea cycle pathway in SIRT5 KO mouse livers.

The urea cycle is a critical biochemical process for ammonia detoxification by converting the toxic ammonia to less toxic urea for excretion via the kidney. Deficiency in urea cycle enzymes or liver failure causes hyperammonemia, and leads to hepatic encephalopathy and life-threatening conditions^25^. In the present study, we observed that 21-month-old SIRT5 KO mice had a higher blood ammonia level than WT mice, and that SIRT5 KO mice had lower tolerance of HAD feeding than WT mice. Interestingly, through behavioral tests, we found that balance and coordination, locomotor activity, and spatial exploration were improved in WT male mice after 4 weeks of HAD feeding. However, due to reduced ammonia tolerance upon HAD feeding, the improvement in coordination and exploratory activity was deprived in SIRT5 KO male mice. Hence, SIRT5 is important in maintaining ammonia tolerance, and preventing ammonia-related neuronal dysfunction.

In the present study, we demonstrated that lysine succinylation regulates the urea cycle in mouse liver. Meanwhile, there are some caveats worth mentioning. First, the way we quantified succinylation stoichiometry (L/(L + H) ×100%) was based on the assumption that succinylation sites are not or negligibly modified by other types of modification, so that endogenously non-succinylation sites can be all chemically succinylated by deuterated succinic anhydride. However, lysines are subject to many different PTMs, and a subset of succinylation sites were reported to overlap with acetylation sites^4,5^. So, the succinylation stoichiometries of some lysine sites may have been at least slightly overestimated in this study due to any undetected discrepancy between the unmodified and total population sizes for that protein lysine site. Second, in addition to desuccinylase activity, SIRT5 has several other deacylase activities, including demalonylase, deglutarylase, and very weak deacetylase, activities. In this study, we demonstrated the importance of the succinylation of ASS1 in mediating SIRT5’s protective effect on urea cycle function. However, another urea cycle enzyme, carbamoyl-phosphate synthase I, has been identified as a deacetylation and deglutarylation substrate of SIRT5^29,30^. Although the stoichiometries of these acylations are unknown, their contributions to a reduced urea cycle function in SIRT5 KO mice is not ruled out.

This study, for the first time, quantified succinylation stoichiometry at a proteome scale in mouse liver and showcased the value of stoichiometry quantification in informing functional importance of succinylation. The finding that succinylation and SIRT5 regulate urea cycle function and ammonia detoxification is interesting because it foreshadows a linkage among succinyl-CoA metabolism, NAD availability, chronic liver disease, and ammonia neurotoxicity.

## Acknowledgements

This project was supported by NIH grant R24DK085610 (E.V. and B.S.), Gladstone Institutes intramural funds (E.V.) and Buck Institute intramural funds (E.V. and B.S.). R.Z. was supported by a postdoctoral fellowship from the Glenn Foundation for Medical Research. X.X. was supported by the NIH (T32GM8806). C.C. was supported by the by a postdoctoral fellowship from the Larry L. Hillblom Foundation. J.G.M. was supported by the NIH (T32 AG000266). B.S. was supported by NIH grant U01AG060906 and NIH Shared Instrumentation Grant 1S10OD016281 (Buck Institute). We thank C. Ronald Kahn, Christopher B. Newgard, and Matthew Hirschey for helpful discussions. We thank Sofiya Galkina for assisting with grip strength tests.

## Author contributions

R.Z., C.C., B.S., and E.V. conceived the study. R.Z., X.X., C.C., B.S., and E.V. contributed to the design of the study and wrote the manuscript with help of the other co-authors. X.X., J.G.M., L.W., J.B. and J.R performed the proteomics analyses, and B.S. supervised proteomics analyses. R.Z., R.K., R.R., and W.H. contributed to mouse maintenance and breeding, performing behavioral tests, and collected mouse tissues and blood. Y.N. prepared samples for metabolomics analyses, X.L. performed metabolomics experiments, and J.W.L. supervised metabolomics analyses. R.Z. constructed and purified recombinant proteins, and measured protein thermal stability and enzymatic activity kinetics. R.Z. performed experiments using cell culture, and measured ammonia levels. C.C., X.X., and R.Z. contributed to the analyses and visualization of proteomics. R.Z. contributed to the analyses and visualization of metabolomics datasets. All authors reviewed the manuscript.

## Supplementary figure legends

**Fig. S1.**
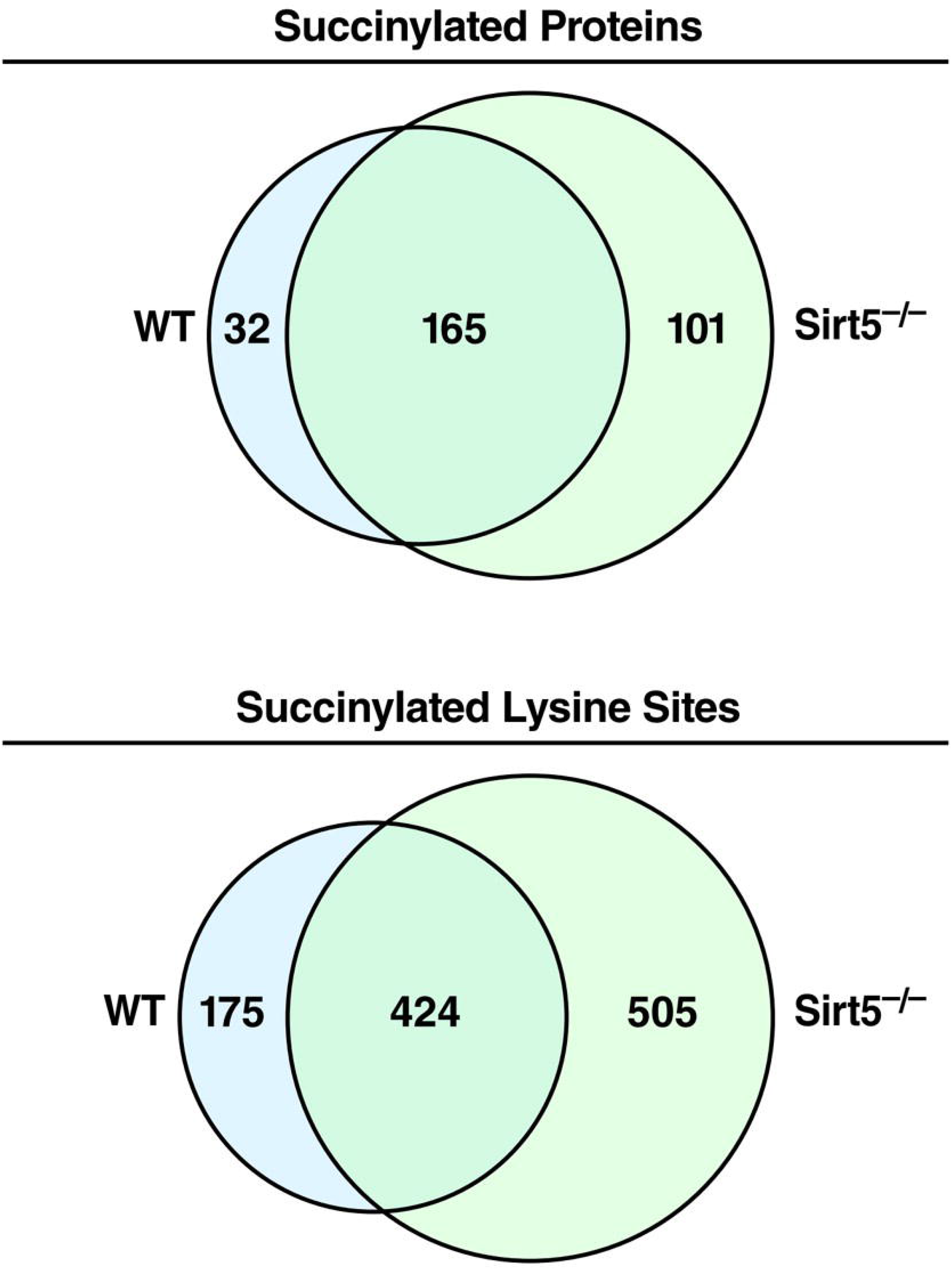
Identification of lysine succinylated proteins and peptides. Venn diagrams showing the overlap of identified succinylated proteins and peptides in WT and *Sirt5*^*-/-*^ mouse liver.

**Fig. S2.**
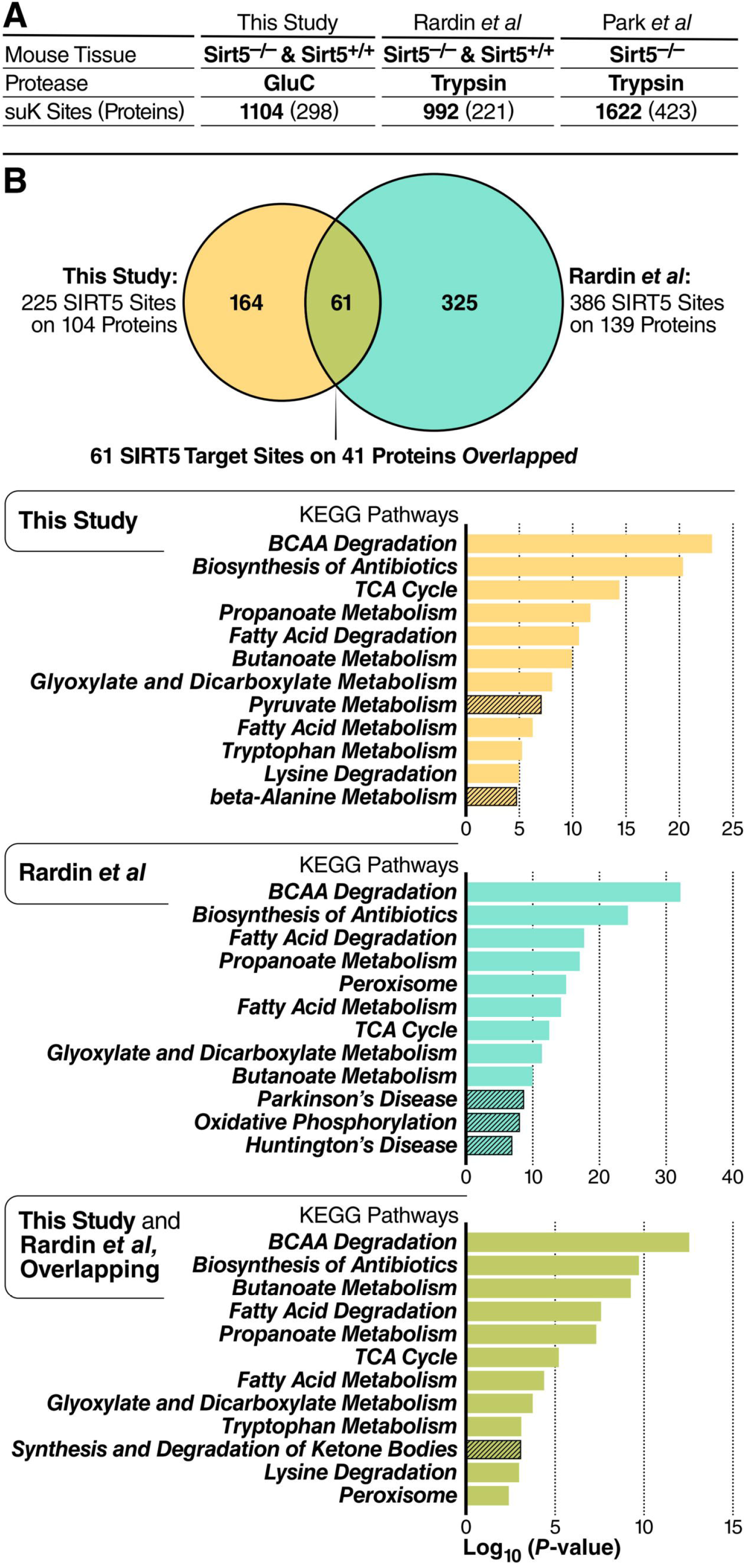
Comparison of immunoenriched succinylomes. **(a)** Comparison of immunoenriched succinylation proteomes performed in this and previous studies (Rardin et al., *Cell Metab*. 2013; Park et al., *Mol Cell*. 2013) ^3,4^. **(b)** KEGG pathway analysis of succinylated proteins identified in this study (yellow circle) or in Rardin et al. (blue circle) or in both studies (green section) that were targeted by SIRT5 (> twofold increase and p < 0.01).

**Fig. S3.**
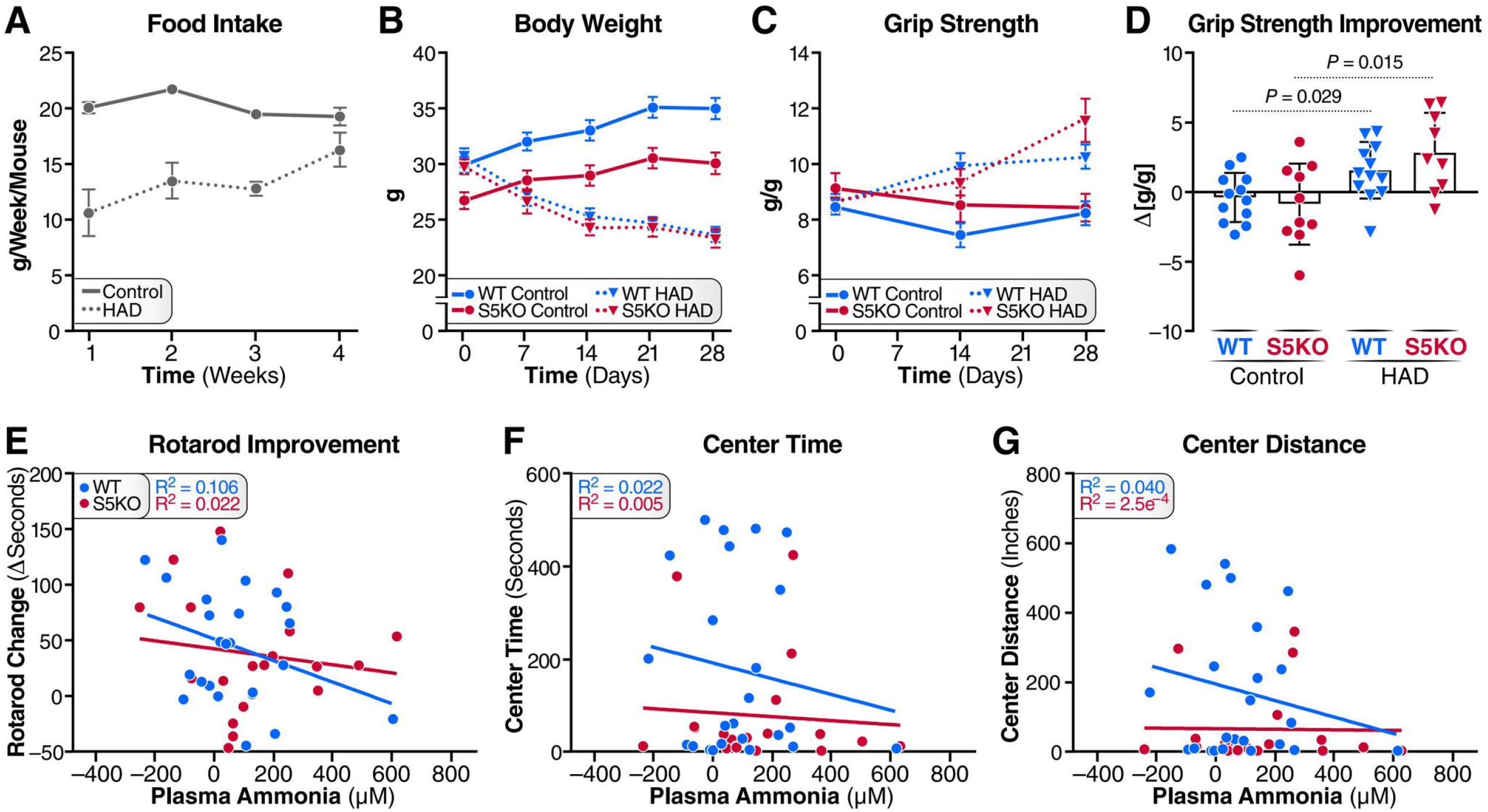
SIRT5 deficiency reduces locomotor activity upon high-ammonium diet feeding in male mice. **(a)** Food intake was recorded weekly per cage and then divided by the number of mice in the cage to calculate the food intake per mouse. n = 4–6. Error bars indicate mean ± s.e.m.. **(b)** Body weight was monitored weekly. n = 9–12. Error bars indicate mean ± s.e.m.. **(c-d)** Grip strength was measured on Days 0, 14, and 28, respectively. Grip strength values were normalized with body weights at the corresponding time points (c). Grip strength improvement was calculated as ΔGripStrength (Day 27–Day 0). n = 9–12. Error bars indicate mean ± s.e.m. P values were generated with unpaired student t test. **(e-g)** Open-field test of WT and *Sirt5*^*-/-*^ male mice was performed after 4 weeks of control diet or HAD feeding. Correlation between rotarod performance improvement and plasma ammonia level (**e**), correlation between center time and plasma ammonia level (**f**), and correlation between center distance and plasma ammonia level (**g**), in WT and S5KO male mice fed on either control diet or HAD, were tested via linear regression. n = 9–12.

