## Supplementary material for "Regulation of urea cycle by reversible high stoichiometry lysine succinylation": Materials and Methods

**Animals.**

Animal studies were performed according to protocols approved by IACUC (the Institutional Animal Care and Use Committee). Wild-type (WT) and *Sirt5*^-/-^ (SIRT5 KO) mice on 129 background were maintained on a standard chow diet (Teklad Global 18% Protein Rodent Diet, ENVIGO, Cat.#2018) until they were put on a special diet or euthanized. For the high-ammonium diet (HAD) feeding model, 12-week old WT and SIRT5 KO male mice were fed on OpenStandard Diet/control chow diet (Research Diets, Inc., D11112201) or HAD (Research Diets, Inc., D19070205) for four weeks. HAD was formulated by adding 20% (wt/wt) ammonium acetate (VWR Chemicals, Cat.#VWRC0103) with OpenStandard Diet. Mice had free access to food and water. Mouse body weight and food intake were monitored weekly. Behavioral tests were performed at certain time points as detailed below.

**Behavioral tests**

1) **Grip strength test**

The grip strength of mouse forelimb and hind limb was measured on Day 0, Day 13, and Day 27 of a special or matched cohort control diet. Grip strength test was performed by the same person throughout this study in a double-blinded manner. Before any measurements were taken, the mouse was lowered over the grid to place its torso parallel with the grid and both its forepaws and hind paws in grid contact. Each mouse was progressively pulled back by its tail while ensuring its torso remained parallel with the grid, and its maximal grip strength value was recorded (BIOSEB, Cat.#BIO-GS3).

2) **Rotarod test**

Rotarod performance (Ugo Basile, Cat.#47750) was measured on Day 1, Day 14, and Day 28 of a special or matched cohort control diet. Mice were habituated for 30 min in the test room before testing. Each test includes one training session and three trial sessions. Consistent white noise (60dB) was displayed throughout habituation and testing sessions to reduce sound interruption. Training sessions comprised the mice staying on the rotating rod for 5 min at a constant speed of 5 rpm. The mouse was gently placed in a separate chamber (up to 5 mice can fit on the rod) before rotation was initiated. The test was performed by doing 3 trials, with breaks of 30 min between each trial. Each Trial session consisted of 5 seconds of constant slow rotation, with rotating rod speed increased gradually over the course of 6 minutes from 5 to 50 rpm. The Trial ends either automatically if the animal falls from the rod as detected by an infrared light sensor, or manually in cases when the mouse gripped the rod and started rotating with it. At the end of the 6 minutes the timer is stopped.

3) **Open-field test**

Open-field test was performed after four weeks of special diet feeding with TRU SCAN Activity Monitoring System (Coulbourn Instruments). Mice were habituated for 30 min in the test room with lights off before testing. Mice were gently placed in a corner of the unit and allowed to move freely for 15 min, while being monitored by the automated tracking system. Tru Scan's precision is 32 × 32, twice the cage's beam resolution; 0.5 inch (1.27 cm) for the beam spacing of the large cage and 0.3 inch (0.76 cm) for spacing of the small cage. The total sum of elapsed time of all movements in the floor plane and the total sum of all vectored coordinate changes (move distance) in the floor plane were calculated. Stereotypic movement is defined as the total number of coordinate changes less than ± 0.999 beam spaces in each floor plane (X and Y) dimension and back to the original point that do not exceed 2 seconds apart. Movements that are less stereotypic were calculated as ambulatory movements. The arena-center is the region that is more than 2.5-beam spaces away from the arena walls.

**Immunoaffinity enrichment succinyl proteomics.**

1) **Materials.**

HPLC solvents, including acetonitrile and water, were obtained from Burdick & Jackson (Muskegon, MI). Reagents for protein chemistry, including triethylammonium bicarbonate buffer (TEAB), iodoacetamide (IAA), dithiothreitol (DTT), formic acid, trifluoroacetic acid (TFA), trichostatin A, urea, nicotinamide, and bovine serum albumin (BSA), were purchased from Sigma Aldrich (St. Louis, MO). Tris(2-carboxyethyl)phosphine (TCEP), and protease/phosphatase inhibitor cocktail were purchased from Thermo Scientific (Rockford, IL), and HLB Oasis SPE cartridges were purchased from Waters (Milford, MA). Sequencing grade trypsin was from Promega (Madison WI). Glu-C endoproteinase was purchased from Roche (Indianapolis, IN). Antibody beads for affinity enrichment was from Cell Signaling Technology (Cat.#13764, PTMScan Succinyl-Lysine Motif [Succ-K] Kit).

2) **Sample preparation of mouse liver for MS succinyl analysis.**

Sample preparation has been described previously^1^. Briefly, 1 mg of protein per mouse was denatured with 8 M urea, 50 mM TEAB, 1 µM trichostatin A, 20 mM nicotinamide, 75 mM sodium chloride, 1x protease/phosphatase inhibitor cocktail (PIC) with HPLC-grade water. Reduce proteins in 4.5 mM DTT at 37 ℃ for 30 min with agitation at 1,400 rpm. Subsequently, alkylate proteins in 10 mM iodoacetamide with incubation in the dark at room temperature for 30 min, and incubated overnight at 37 ℃ with sequencing grade Glu-C added at a 1:50 enzyme:substrate ratio (wt/wt). Samples were then acidified with formic acid and desalted using HLB Oasis SPE cartridges. Samples were eluted, concentrated to near dryness by vacuum centrifugation, and resuspended in cold 1x immuno-affinity purification (IAP) buffer. Incubate peptides with succinyl-lysine antibody beads according to manufacturer’s instructions at 4 ℃ overnight with agitation. Beads were washed twice in IAP buffer, and three times in HPLC-grade water. The peptides were eluted by washing twice in 0.15% TFA. The succinyllysine peptide enrichments were subsequently desalted using C18 ZipTips. After evaporation of organic solvents, samples were suspended in 0.2% formic acid and analyzed by LC-MS/MS on a TripleTOF 6600 mass spectrometer. A solution-only “blank” or auto-calibration was examined between sample acquisitions to prevent carry over that would affect downstream quantitative analysis.

3) **Data acquisition using DDA and DIA.**

Data-dependent acquisitions (DDA) were used to create a custom spectral library and data-independent acquisitions (DIA) were used for quantification. Samples were analyzed by reverse-phase HPLC-ESI-MS/MS using an Eksigent Ultra Plus nano-LC 2D HPLC system (Dublin, CA) combined with a cHiPLC system coupled to a SCIEX TripleTOF 6600 mass spectrometer (SCIEX, Redwood City, CA). After injection, peptide mixtures were transferred onto a C18 pre-column chip (200 μm × 6 mm ChromXP C18-CL chip, 3 μm, 300 Å, SCIEX) and desalted the peptides by washing with mobile phase A at 2 µL/min for 10 min. Then, peptides were transferred to an analytical column (75 μm × 15 cm ChromXP C18-CL chip, 3 μm, 300 Å, SCIEX), and eluted at a flow rate of 300 nL/min with a 2-3 h gradient using mobile phases A and B. Specifically, use a linear gradient from 5% mobile phase B to 35% mobile phase B over 80 min. Subsequently, ramp the mobile phase B to 80% over 5 min, then hold at 80% B for 8 min before returning to 5% B for a 25 min re-equilibration.

Building a MS instrument method for DDA: MS1 precursor ion scan from m/z 400-1,500 (accumulation time of 250 ms). Set the intensity threshold to trigger MS/MS scans for ions of charge states 2-5 to 200 counts. Set the dynamic exclusion of precursor ions to 60 s. MS/MS product ion scan with a MS2 scan range from m/z 100-1,500 (accumulation time of 100 ms per each 30-product ion scan per cycle)^2^. Set the collision energy spread to CES = 5, then select the "high sensitivity product ion scan mode".

Building a MS instrument method for DIA: Perform MS1 precursor ion scan from m/z 400-1,250 (accumulation time of 250 ms), and MS/MS product ion scans for 64 variable SWATH segments with a MS2 scan range from m/z 100-1,250 (accumulation time of 45 ms per each 64-product ion scan per cycle)^3,4^. Set the collision energy spread to CES = 10, then select the "high sensitivity product ion scan mode". Use the 64 variable window DIA/SWATH acquisition strategy as previously described^3^ to obtain label-free quantification with a total cycle time of ~3.2 s.

4) **Data analysis.**

MS database search engine (ProteinPilot 5.0, SCIEX, Framingham, MA) was used to analyze DDA acquisitions. For the database searches, the enzyme was set to GluC, Mus musculus was set for the species, fixed medications were set as carbamidomethylation for cysteine, and methionine oxidation (+15.99), glutamate to pyroglutamate conversion (-17.02), and succinylation (+100.01) were defined as variable modifications. False discovery rate (FDR) was set up to be < 1%. Identification confidence thresholds per peptide were set as 99. The search engine results from ProteinPilot were subsequently imported into Skyline to build spectral libraries for SWATH data processing with the DIA acquisition data in Skyline (B. MacLean et al., 2010, Bioinformatics), an open-source software project (http://proteome.gs.washington.edu/software/skyline). In Skyline, the XICs of the top 3 precursor ions (e.g., M, M+1 and M+2) were extracted and quantitative SWATH MS2 data analysis was based on extracted ion chromatograms (XICs) of up to 10 of the most abundant fragment ions in the identified spectra. Extracted peak areas were summed per target peptide/protein.

**Quantification of Lysine Succinylation Stoichiometry.**

1) **Chemicals.**

Acetonitrile and water were obtained from Burdick and Jackson (Muskegon, MI, USA). Reagents for protein chemistry including iodoacetamide, dithiothreitol (DTT), ammonium bicarbonate, formic acid (FA), urea, and succinic anhydride-2,2′,3,3′-d4 (>98% deuterium atom enrichment) were purchased from Sigma Aldrich (St. Louis, MO, USA). BSA and acetylated BSA were purchased from Pierce (Rockford, IL, USA). Sequencing grade Glu-C endoproteinase was purchased from Roche (Indianapolis, IN, USA). HLB Oasis SPE cartridges were purchased from Waters (Milford, MA, USA).

2) **Chemical treatments to quantitatively succinylate proteins.**

Protein lysates from mouse livers were prepared as described above. Per-succinylation: 100 μg of protein was diluted to 1 μg/uL using ammonium bicarbonate solution (8 M urea, 200 mM ammonium bicarbonate pH 8). Dithiothreitol was added to a final concentration of 20 mM and incubated at 37°C for 30 min. Iodoacetamide was added to a final concentration of 40 mM and incubated in the dark at room temperature for 30 min. The succinic anhydride-d4 powder was dissolved in dry DMSO prior to the reaction. Chemical per-succinylation was performed as previously described^5-7^ for per-acetylation and subsequently adopted for succinylation analysis according to a protocol from Meyer et al.^5^ by adding 60 μmol of succinic anhydride-d4 to the sample and incubating in a Thermomixer at 4 °C for 20 min. The chemical succinylation process was repeated twice more. Ammonium hydroxide was added between each per-succinylation reaction to increase the pH to ~8, and 5 μL of 50% hydroxylamine was added after the final reaction to revert O-succinylation side reactions (note: the labeling reaction can also be carried out using 200 mM triethanolamine bicarbonate, TEAB, instead of ammonium bicarbonate). The urea concentration was diluted to 0.8 M using 25 mM ammonium bicarbonate and the protease Glu-C was added at a 1:50 ratio (wt:wt) and incubated overnight at 37 °C. Samples were then acidified by the addition of formic acid to 1% by volume and desalted using Oasis HLB extraction cartridges. The eluate was dried in a Speed-Vac and resuspended in 0.1% formic acid with 2% ACN.

3) **Offline peptide fractionation by basic-pH reversed phase HPLC.**

Per-succinylated and Glu-C digested proteins from whole liver lysates were separated by a Waters 1525 binary HPLC pump system including a Waters 2487 UV detector. The high-pH reversed phase peptide separation was performed using an Agilent Zorbax 300Extend C18 column (4.6 mm × 250 mm, 5 μm particle size; Agilent, Santa Clara, CA, USA). Peptides were separated at a flowrate of 0.7 mL/min using the following gradient: 100% A to 92% A over 7.3 min, 92% A to 73% A over 38 min, 73% A to 69% A over 4 min, 69% A to 61% A over 16 min, 61% A to 40% A over 7 min, 40% A to 10% A over 7.73 min, constant 10% A from 80 to 85 min, 10% A to 100% from 85 to 86 min, and finally re-equilibrated in 100% A for 24 min. Buffer A was 10 mM ammonium formate in water, and buffer B was 10 mM ammonium formate in 90% ACN and 10% water. The pH of both mobile phases adjusted to 10 with neat ammonia. Fractions were automatically collected and every eighth fraction was pooled, resulting in eight final fractions.

4) **Mass Spectrometry.**

Samples were analyzed by reverse-phase HPLC-ESI-MS/MS using the Eksigent Ultra Plus nano-LC 2D HPLC system (Dublin, CA, USA) combined with a cHiPLC System, which was directly connected to a quadrupole time-of-flight SCIEX TripleTOF 6600 mass spectrometer (SCIEX, Redwood City, CA, USA). Typically, mass resolution in precursor scans was 45,000 (TripleTOF 6600), whereas fragment ion resolution was ~15,000 in ‘high sensitivity’ product ion scan mode. After injection, peptide mixtures were transferred onto a C18 precolumn chip (200 μm × 6 mm ChromXP C18-CL chip, 3 μm, 300 Å, SCIEX) and washed at 2 μL/min for 10 min with the loading solvent (H2O/0.1% formic acid) for desalting. Subsequently, peptides were transferred to the 75 μm × 15 cm ChromXP C18-CL chip, 3 μm, 300 Å, (SCIEX), and eluted at a flow rate of 300 nL/min using a 2 h gradient using aqueous and acetonitrile solvent buffers.

Some initial data-dependent acquisitions (DDA) were carried out to obtain MS/MS spectra for the 30 most abundant precursor ions (100 ms per MS/MS) following each survey MS1 scan (250 ms), yielding a total cycle time of 3.3 s as previously described^8^. For collision induced dissociation tandem mass spectrometry (CID-MS/MS), the mass window for precursor ion selection of the quadrupole mass analyzer was set to ±1 m/z using the Analyst 1.7 (build 96) software. All stoichiometry study samples were analyzed by data independent acquisitions (DIA), or specifically variable window SWATH acquisitions. In these SWATH acquisitions, instead of the Q1 quadrupole transmitting a narrow mass range through to the collision cell, windows of variable width (5 to 90 m/z) are passed in incremental steps over the full mass range (m/z 400–1250). The cycle time of 3.2 s includes a 250 ms precursor ion scan followed by 45 ms accumulation time for each of the 64 SWATH segments. The variable windows were determined according to the complexity of the typical MS1 ion current observed within a certain m/z range using a SCIEX ‘variable window calculator’ algorithm (i.e., more narrow windows were chosen in ‘busy’ m/z ranges, wide windows in m/z ranges with few eluting precursor ions).

5) **Data processing and bioinformatics**.

Mass spectrometric wiff files were first converted to mzML using the SCIEX data converter version 1.3, and subsequently mzML files were converted to mzXML files using ProteoWizard version 3.0.8851. The DIA-Umpire^9^ signal extraction module was used to process SWATH acquisitions, which detects correlated precursor and fragment ion features and assembles them into pseudo-tandem MS/MS spectra stored in mgf files. These pseudo MS/MS spectra derived from all SWATH acquisitions were then searched using the database search engine Mascot^10^ server version 2.3.02. In addition, DIA-Umpire-derived mgf files containing pseudo MS/MS spectra were searched with MSGF+^11^, refined with PeptideProphet^12^, and combined with iProphet^13^. For the database searches, enzymatic cleavage sites were defined at Glu (E) and Asp (D), and fixed modifications were set as carbamidomethylation for cysteine residues. Variable modifications were defined for lysine residues as light and heavy succinyl for succinylation occupancy experiments. False discovery rates (FDR) were required to be <0.01. The search engine results were subsequently used in Skyline to build spectral libraries for SWATH data processing in Skyline. SWATH acquisitions were quantitatively processed using Skyline 3.5^14^, an open source software project (http:// proteome.gs.washington.edu/software/skyline). Quantitative SWATH MS2 data analysis was based on extracted ion chromatograms (XICs) of up to 10 of the most abundant fragment ions in the identified spectra, but in contrast to MS1 quantification, only ions containing the modified lysine of interest were used for quantification, as those can be used to differentiate between the light and heavy acylated peptide forms. Occupancy measurements were computed from light (L) and heavy (H) peak areas as L/(L + H).

For peptides containing only one lysine, Python (version 2.7.10) scripts written in-house were used to determine the differentiating fragment ions and compute the stoichiometry from custom Skyline reports. To enable quantification of peptides containing more than one lysine modification, an additional R script (http://www.R-project.org/), referred to as StoichiolyzeR (<https://github.com/GibsonLab/StoichiolyzeR>), was written in-house and uses the package mzR^15^ to extract fragment ion signal from the mzXML file and to compute the stoichiometry of acylation sites. Briefly, the StoichiolyzeR program requires an mzXML file generated from the original wiff raw data file, a text file containing the SWATH isolation window definitions, and a text file containing peptide information, such as a Skyline report. The peptide information text file contains the peptides sequence, peak retention time borders for peak area extraction/integration, precursor charge, mzXML file name, and the peptide sequence residue numbers to define the K-succinyl sites within the protein. The program first determines all theoretical heavy and light fragment ion m/z values that contain the lysine residue(s). The program also confirms that the heavy/light precursor ion pairs were sampled in the same SWATH isolation window. Next, the heavy and light fragment peak areas are extracted using a predefined ppm mass tolerance within the given retention time elution range. Fragment ion peak areas subsequently were filtered by a user-defined minimum heavy peak area to avoid reporting very noisy, low abundant ions; however, if heavy peak areas are below the set threshold, the same threshold is tested for the corresponding light fragment ion (in case a specific K-succinyl site shows a very high stoichiometry in which case heavy peak areas could be very low). Fragment ions are further processed if either the heavy or the light peak area is above the threshold. The ratios of light/(light + heavy) are computed, and both the median ratio and the ratio from the highest ranked (differentiating) fragment ion (rank 1) are reported. The stoichiometry for peptides containing two lysine residues per peptide is computed for the first and second lysine using the differentiating b-ion and y-ion ratios, respectively.

**Data availability:** Raw data and complete MS data sets have been uploaded to the Center for Computational Mass Spectrometry, to the MassIVE repository at UCSD, and can be downloaded using the following link:

http://massive.ucsd.edu/ProteoSAFe/status.jsp?task=10c82e20ad4d4231b3f762d3fa40af81

The mass spectrometric raw data are deposited with the MassIVE ID number MSV000089726; it is also available at ProteomeXchange with the ID PXD034880.

[Note to the reviewers: To access the data repository MassIVE (UCSD) for MS data, please use: Username: MSV000089726_reviewer; Password: winter].

**Expression and purification of recombinant ASS1 proteins.**

His-tagged human ASS1 plasmid was a gift from Nicola Burgess-Brown (Addgene plasmid #42368; <http://n2t.net/addgene:42368>; RRID:Addgene_42368). Mutant ASS1 plasmids were constructed with a site-direct mutagenesis kit (New England Biolabs, Cat.#E0554) according to manufacturer’s instructions. WT and mutant ASS1 proteins were expressed using BL21(DE3) competent cells. Briefly, transformed bacteria were plated on lysogeny broth agar plates with 50 mg/mL kanamycin, and incubate overnight at 37°C. Resuspend a single colony in 5 ml liquid culture with Kanamycin. Culture at 37°C for 16 hr. Inoculate 4 mL culture to 200 mL LB media containing kanamycin. Measure OD600. When OD600 reaches 0.6, equilibrate culture at 30°C for 1 hr. The expression of recombinant proteins was induced with 0.5 mM IPTG in LB media containing Kanamycin. Culture at 30°C for 16 hr.

Centrifuge culture at 4°C to collect pellet. Lyse bacteria collected from 200 mL media with 8 mL lysis buffer (50 mM sodium phosphate dibasic, pH8.0,	300 mM NaCl, 10 mM imidazole, 1× EDTA-free protease inhibitors,	75 U/mL benzonase, 1 mg/mL lysozyme). The recombinant proteins in the cleared lysates were purified using HisPur Cobalt Purification Kit, 0.2 mL (Thermo Scientific, Cat.#90090) according to manufacturer’s instructions. Imidazole was removed with a desalting buffer (20 mM Tris-HCl, pH 7.8, and 0.25 M NaCl, 1 mM DTT, 10% glycerol) via centrifugation with 30 kDa centrifugal filter for 4 times. Protein concentration was measured with Branford reagent at A595.

**Differential scanning fluorimetry**

To monitor thermal unfolding of recombinant WT and mutant ASS1 proteins, differential scanning fluorimetry using a thermal shift protein stability kit (Biotium, Cat.#33021) according to manufacturer’s instructions. The enzymes were heated from 25.0°C to 99.0°C with a ramp rate of 0.5 °C/min in a CFX96 Touch Real-Time PCR Detection System (BioRad). The obtained fluorescence was normalized so that minimum value was set to 0 and maximum set to 1. The temperature at which 50% of protein is melted (Tm, in °C) was determined by Boltzmann sigmoidal analysis after curve fitting using GraphPad Prism 6 (GraphPad Software).

**ASS1 activity assay.**

ASS activity was determined based on accumulation of the product pyrophosphate as inorganic phosphate, following cleavage with pyrophosphatase as previously described^16^. The kinetics of enzyme activity was determined as previously described^17^. Briefly, 0.5 mg/mL of enzyme were incubated for 30 min at 37°C in an assay mixture (20 mM HEPES, pH7.0, 1 mM L-aspartate, 1 mM L-citrulline, 2 mM ATP, 5 mM MgCl_2_) with serially diluted substrates (L-aspartate or l-citrulline). When the concentration of a substrate was varied, the other substrate was kept at 1 mM concentration. Incubate reaction solutions at 37°C for 30 min. Add equal volume of HPLC grade methanol, mix, let sit on ice for 5 min. Centrifuge at 15000 g at 4°C for 10 min. The production of pyrophosphate in the supernatants was analyzed using a pyrophosphate assay kit (Sigma-Aldrich, Cat.#MAK168) according to manufacturer’s instructions. To determine Km and Vmax, data were fitted by non-linear regression to the Michaelis-Menten equation. Each data point is the average of three separate measurements.

**Metabolomics analysis.**

1) **Sample preparation for metabolomics analysis.**

Intracellular metabolites from mouse liver tissues (5-10 mg) were extracted using 80% methanol/water as the extraction solvent ^18-20^. After extraction, metabolite extract in 80% methanol/water was split into two tubes (one for polar metabolite analysis and the other one for acyl-CoA analysis), and then extraction solvent was evaporated using a speed vacuum concentrator. Dry pellets were stored in −80 °C freezer until ready for LC-MS analysis. For acyl-CoA analysis, dry pellets were reconstituted into 30 μL of sample solvent (water containing 50 mM ammonium acetate) per 6 mg tissue, and 8 μL was injected into the LC-MS. For non-acyl-CoA polar metabolite analysis, pellets were reconstituted into 30 μL of sample solvent (water:methanol:acetonitrile, 2:1:1, v/v/v) per 3 mg tissue, and 3 μL was injected into the LC-MS.

2) **LC/MS and data analysis.**

A Dionex 3000 UHPLC was used for metabolite separation. For polar metabolite analysis, a hydrophilic interaction chromatography method (HILIC) with an Xbridge amide column (100 × 2.1 mm i.d., 3.5 μm; Waters) was used, and detailed parameter settings were according to methods described in previous studies ^21^. For acyl-CoA analysis, a reversed phase liquid chromatography method employing a Luna C18 column (100 × 2.0 mm i.d., 3 μm; Phenomenex) was used ^20^. LC was coupled with Q Exactive mass spectrometer (Thermo Scientific) for metabolite separation and subsequent detection. The Q Exactive mass spectrometer was equipped with a HESI probe, and for non-acyl-CoA polar metabolite analysis, it was operated in the full-scan mode with positive/negative switching with the resolution set at 70 000 (at m/z 200). When it was used for acyl-CoA analysis, it was operated in the positive ion mode with a full-scan range of 150–1000 (m/z).

LC-MS data were analyzed using Sieve 2.0 (Thermo Scientific), and the integrated area under metabolite peak was used to compare the relative abundance of each metabolite in different samples in the same batch. Retention times for each metabolite are based on in-house library and determined by comparing to standard compounds when available, m/z and characteristic fragments. When multiple metabolites with the same m/z cannot be easily distinguished from each other, then they are combined into one row and the chemical formula is used to represent the mixed isomers.

**Data availability:** Metabolomics data for WT (*Sirt5^+/+^*) and *Sirt5^-/-^* mouse livers were deposited at Metabolomics Workbench (Project ID: PR001365. Project DOI: 10.21228/M88Q5H).

**Ammonia measurement.**

Ammonia levels in cell culture media and mouse blood sera were measured using an ammonia assay kit (Sigma-Aldrich, Cat.#AA0100) according to manufacturer’s instructions. Plasma samples for ammonia measurement were prepared from 21 months old *Sirt5*^-/-^ (S5KO), heterozygous (Het), WT mice after 24 hr fasting; and from 4 months old S5KO and WT mice fed on OpenStandard Diet or HAD without fasting. Plasma samples were centrifuged at 13000 g and 4°C for 10 min to remove any insoluble precipitate before measurement.
